# Common molecular features of H3K27M DMGs and PFA ependymomas map to hindbrain developmental pathways

**DOI:** 10.1101/2022.11.17.516833

**Authors:** Matthew Pun, Drew Pratt, Patricia R. Nano, Piyush K. Joshi, Li Jiang, Bernhard Englinger, Arvind Rao, Marcin Cieslik, Arul M. Chinnaiyan, Kenneth Aldape, Stefan Pfister, Mariella G. Filbin, Aparna Bhaduri, Sriram Venneti

**Affiliations:** Laboratory of Brain tumor metabolism and epigenetics, Department of Pathology, University of Michigan, Ann Arbor, MI, USA.; Chad Carr Pediatric Tumor Center, Department of Pediatrics, University of Michigan, Ann Arbor, MI, USA.; Cellular and Molecular Biology Program, University of Michigan Medical School, Ann Arbor, MI 48109, USA.; Medical Scientist Training Program, University of Michigan Medical School, Ann Arbor, MI 48109, USA.; Laboratory of Pathology, Center for Cancer Research, National Cancer Institute, National Institutes of Health, 10 Center Dr., Room 2S235, Bethesda, MD, 20892, USA.; Department of Biological Chemistry, University of California, Los Angeles, Los Angeles, CA 90095, USA.; Hopp Children’s Cancer Center (KiTZ) Heidelberg, Division of Pediatric Neurooncology, German Cancer Consortium (DKTK), German Cancer Research Center (DKFZ), 69120 Heidelberg, Germany.; Department of Pediatric Oncology, Dana-Farber Boston Children’s Cancer and Blood Disorders Center, Boston, MA 02115, USA.; Broad Institute of Harvard and MIT, Cambridge, MA 02142, USA.; Department of Computational Medicine and Bioinformatics, University of Michigan Medical School, Ann Arbor, MI 48109, USA.; Department of Biostatistics, University of Michigan, Ann Arbor, MI 48109, USA.; Department of Radiation Oncology, University of Michigan Medical School, Ann Arbor, MI 48109, USA.; Michigan Center for Translational Pathology, Department of Pathology, University of Michigan Medical School, Ann Arbor, MI 48109, USA.; Rogel Cancer Center, University of Michigan Medical School, Ann Arbor, MI 48109, USA.; Department of Urology, University of Michigan Medical School, Ann Arbor, MI 48109, USA.; Howard Hughes Medical Institute, University of Michigan Medical School, Ann Arbor, MI 48109, USA.; Division of Pediatric Neurooncology, German Cancer Research Center (DKFZ) and German Cancer Consortium (DKTK), Heidelberg 69120, Germany.; Department of Pediatric Hematology and Oncology, Heidelberg University Hospital, Heidelberg 69120, Germany.

**Keywords:** cancer, chromatin biology, onco-histones, pediatric tumors, brain development, neuro-oncology

## Abstract

Globally decreased histone 3, lysine 27 tri-methylation (H3K27me3) is a hallmark of H3K27-altered diffuse midline gliomas (DMGs) and group-A posterior fossa ependymomas (PFAs). H3K27-altered DMGs are largely characterized by lysine-to-methionine mutations in histone 3 at position 27 (H3K27M). Most PFAs overexpress EZH inhibitory protein (EZHIP), which possesses a region of similarity to the mutant H3K27M. Both H3K27M and EZHIP inhibit function of the polycomb repressive complex 2 (PRC2) responsible for H3K27me3 deposition. These tumors often arise in neighboring regions of the brainstem and posterior fossa. In rare cases PFAs harbor H3K27M mutations, and DMGs overexpress EZHIP. These findings together raise the possibility that certain cell populations in the developing hindbrain/posterior fossa are especially sensitive to modulation of H3K27me3 states. We identified shared molecular features by comparing genomic, bulk transcriptomic, chromatin-based profiles, and single-cell RNA-sequencing (scRNA-seq) data from the two tumor classes. Our approach demonstrated that 1q gain, a key biomarker in PFAs, is prognostic in H3.1K27M, but not H3.3K27M gliomas. Conversely, Activin A Receptor Type 1 (ACVR1), which is associated with mutations in H3.1K27M gliomas, is overexpressed in a subset of PFAs with poor outcome. Despite diffuse H3K27me3 reduction, previous work shows that both tumors maintain genomic H3K27me3 deposition at select sites. We demonstrate heterogeneity in shared patterns of residual H3K27me3 for both tumors that largely segregated with inferred anatomic tumor origins and progenitor populations of tumor cells. In contrast, analysis of genes linked to H3K27 acetylation (H3K27ac)-marked enhancers showed higher expression in astrocytic-like tumor cells. Finally, common H3K27me3-marked genes mapped closely to expression patterns in the human developing hindbrain. Overall, our data demonstrate developmentally relevant molecular similarities between PFAs and H3K27M DMGs and support the overall hypothesis that deregulated mechanisms of hindbrain development are central to the biology of both tumors.

## Introduction

Advances in high-throughput sequencing have provided incredible windows into understanding the diversity and heterogeneity in cancers. Genomic, transcriptomic, and epigenomic studies have aided in molecular classification of tumors, elucidation of candidate driving events, and potential cells of origin for many cancers, including pediatric brain tumors. Nevertheless, brain tumors remain the leading cause of mortality among all pediatric cancers [17]. Unlike in adult populations, pediatric brain tumors frequently arise in infratentorial regions [62], suggesting a spatiotemporal susceptibility to tumorigenesis. Indeed, transcriptional analysis of pediatric brain tumors including medulloblastoma, gliomas, and ependymomas suggest shared expression programs within certain developmental lineages [15, 20, 22, 25, 29, 35, 69, 73].

Group A posterior fossa ependymomas (PFAs) and diffuse midline gliomas (DMGs), H3K27-altered, are characterized by global reduction of the repressive histone modification histone 3 lysine 27 trimethylation (H3K27me3) [8, 39, 67, 70, 77]. Global H3K27me3 depletion is mediated by inhibition of the function of the polycomb repressive complex-2 (PRC2), which contains the H3K27-specific methyltransferase enhancer of zeste homolog 2 (EZH2). The majority of these DMGs harbor missense mutations in histone 3-encoding genes (primarily *H3-3A* and *H3C2*) [67, 70, 77] that result in a lysine-to-methionine substitution at position 27 (H3K27M). This histone tail mutation inhibits PRC2 activity [8, 14, 39]. Most PFAs, in contrast, exhibit overexpression of EZH inhibitory protein (EZHIP) [4, 57] that similarly suppresses PRC2 activity. EZHIP contains a methionine residue at position 406 that mimics the H3K27M mutant histone [32–34, 61, 65] . In addition to depleting H3K27me3, each of these tumors demonstrates increased global levels of the activation-associated mark H3K27 acetylation (H3K27ac) [33, 36, 53, 57, 59–61, 65].

With their epigenetic similarities, H3K27M DMGs and PFAs also share sensitivities to inhibitors targeting histone-modifying enzymes, including histone lysine demethylases, and histone deacetylases [3, 26, 28, 53, 59, 60]. Although H3K27me3 levels are greatly reduced globally, specific regions of the genome retain H3K27me3, most of which are CpG islands commonly known to be canonical PRC2 binding sites [6, 8, 27, 33, 43]. Moreover, both tumor types appear to rely on residual EZH2 activity. Both pharmacologic and genetic inhibition of EZH2 reduce tumor viability in models of H3K27M DMGs and PFAs [43, 49, 51], suggesting that the tumors rely on residual PRC2 activity.

In addition to their molecular similarities, PFAs and H3K27M DMGs arise in nearby structures associated with the hindbrain. The floor of the fourth ventricle is formed by the pons and the roof by the cerebellum. Whereas the majority of H3K27M DMGs arise from the pons, most PFAs are found in association with the fourth ventricle and cerebellum [44, 58]. Moreover, it is increasingly recognized that a rare population of H3- wildtype, low-H3K27me3 DMGs overexpress EZHIP, and that a similarly small percentage of PFAs harbor H3K27M mutations [11, 46, 57, 64]. The extent to which these two tumor subtypes share biology, and how this relates to hindbrain development, remains an unanswered question. To address this gap in our knowledge, we hypothesized that shared molecular features of H3K27M DMGs and PFAs will reveal key programs within the developing human hindbrain. We addressed this hypothesis by examining genomic, transcriptomic, and epigenomic characterizations of each tumor type to characterize shared biology to better understand the origins and potential vulnerabilities of these tumors.

## Methods

### Copy number alteration analyses

Arm-length copy number alterations for PFAs and DMGs were obtained from supplementary data of published datasets [44, 58]. For tumor samples from the NCI, tumors were classified with the Molecular Neuropathology classifier (v11b6) [10]. Genomic Identification of Significant Targets in Cancer (GISTIC) version 2.0.23 was used to identify and score broad copy number events. Score cutoffs of +/-0.3 were used to call a gain or loss of a chromosome arm consistent with the methods employed by Mackay *et al.* [44].

### Gene expression analyses

PFA (GSE100240, GSE64415) microarray expression data and DMG (Mackay *et al*. 2017) meta-analysis expression data were obtained. A limited differential expression analysis was conducted by dividing the expression datasets by the variable of interest and applying a two-sided student’s t test. Multiple testing correction was applied by the Benjamini-Hochberg method. For gene set enrichment analysis (GSEA), ranked gene lists were generated by taking the product of the negative log-10 transformation of the q-value and the difference in expression for each gene. GSEA 4.2.3 was used to run GSEA Preranked on the resulting gene lists with the following parameters: 15-500 as min-max gene set size; weighted scoring scheme; meandiv normalization; Abs_max_of_probes mode. For standard deviation analyses, the probe with the maximum average value across samples in PFAs was used. Any genes that did not appear in both gene sets were excluded from plotting.

### Histone 3 lysine 27 (H3K27) trimethylation ChIP-seq analyses

For samples from Bender *et al*. [8], BED files with hg19 alignment were obtained from the authors. BED files aligned to hg19 from Harutyunyan *et al*. [27] were obtained from the Genetics and Genomics Analysis Platform (GenAP). Data aligned to hg19 from Bayliss *et al*. [6] (GSE89451) and Mack *et al*. [43] (GSE89451) were obtained from GEO. Bedtools 2.29.2 was used to intersect peaks from each dataset with a custom BED file containing genes and their promoters defined as 2 kb upstream from the transcription start site (TSS). Genes with peaks in greater than two-thirds of tumor samples for each class of tumor were defined as common H3K27me3-retaining genes and used for downstream analyses.

### H3K27ac and enhancer analyses

We obtained published H3K27M DMG-specific enhancer-associated genes from Krug *et al*. [36] and PFA-specific enhancer-associated genes from Mack *et al*. [42]. We computed the overlap in genes from these two gene sets. Each of these curated gene sets was in part derived from comparison against enhancer profiles of other brain tumors. To identify enhancers that were unique to each tumor type independent of these comparisons with non-PFAs and non-H3K27M DMGs, we accessed the ChIP-seq files available from each study and reanalyzed them. We conducted Rank-ordering of Super Enhancer (ROSE) analysis to call super enhancers for each tumor. We examined if PFA-specific enhancers from the original analysis were called as super enhancers in any of the DMG ChIP-seq samples. If an enhancer appeared at least once in the DMG samples, we included it the shared enhancer list. We repeated the converse process for DMG-specific enhancers. This method allowed us to avoid biases generated by comparisons with non-PFAs and non-H3K27M DMGs. Pathway analyses of shared or unique enhancers were completed using Enrichr [16, 38, 79]. Volcano plots were generated using code generated by Appyter.

### Single-cell RNA-seq analyses

Published RNA-Seq by Expectation-Maximization (RSEM) values were obtained for PFA and DMG datasets [20, 25] and loaded as Seurat objects using Seurat (4.1.1) and SeuratObject (4.1.0). Each dataset was independently normalized, and then variable features were selected. Seurat’s SelectIntegrationFeatures and FindIntegrationAnchors were sequentially applied to the datasets, and then IntegrateData was applied based on the anchors identified. The resulting integrated dataset underwent dimensional reduction and clustering. FindConservedMarkers was used to identify differentially expressed genes by cluster. Developing brain single-cell RNA-seq data from Aldinger *et al.* [1] were obtained from https://cbl-dev.cells.ucsc.edu and analyzed using Seurat. AddModuleScore was used to generate a score for the H3K27me3-retained gene signature. Single-cell RNA-seq data from Eze *et al.* [17] was obtained from https://cells-test.gi.ucsc.edu/?ds=early-brain and processed using Seurat as described in [17], and *CRABP1* feature plots were generated using the Seurat ViolinPlot function, ggplot2 (3.3.5), and ggprism (1.0.3). For area under the curve measurements from the Sepp *et al*. [68] dataset, raw count measurements were processed with AUCell (from Bioconductor 3.16) using the H3K27me3-retained gene signature.

### Allen Brain Atlas analyses

Human data were obtained from BrainSpan, and mouse data were obtained from the Allen Developing Mouse Brain map. All resulting analyses were completed with GraphPad Prism 9.1.1.

### Statistical analyses

Graphs were plotted and statistical analyses were performed using Prism software (versions 9.1.1, Graphpad, La Jolla, CA). Data are represented as the means ± standard deviation (S.D.) or as violin plots. The sample size (n) along with the statistical test performed and corresponding p-values are indicated in each figure or figure legend. Progression-free and overall survival metadata, when available, was analyzed comparing survival curves utilized the Log-Rank (Mantel-Cox) test for significance. Unpaired two-tailed, two-sided, Student’s t test or one-way ANOVA followed by multiple comparisons analysis using either Ordinary one-way ANOVA or Kruskal-Wallis tests were used to analyze data. Data were considered significant if p values, adjusted where appropriate, were below 0.05 (95% confidence intervals).

## Results

### H3.1K27M and EZHIP-DMGs share key copy-number alterations with PFAs

To characterize the shared features of H3K27M DMG and PFAs, we began by comparing copy number changes in in two published data sets and from Mackay *et al*. (2017, H3K27M DMG = 295) and Pajtler *et al*. (2015, PFA = 240) [44, 57]. Overall, H3K27M DMGs demonstrated far greater genomic instability compared to PFAs. Nevertheless, comparison of copy number alterations demonstrated that PFAs harbor several recurrent chromosome arm alterations that are shared with H3K27M DMGs. These included gain of the long arm of chromosome 1 (1q) and loss of 6q (**Fig. 1a**). We validated these findings in two independent, non-overlapping tumor cohorts (PFA=573 and H3K27M=271) curated by the National Cancer Institute (NCI) (**Fig. 1b**).

**Fig. 1:**
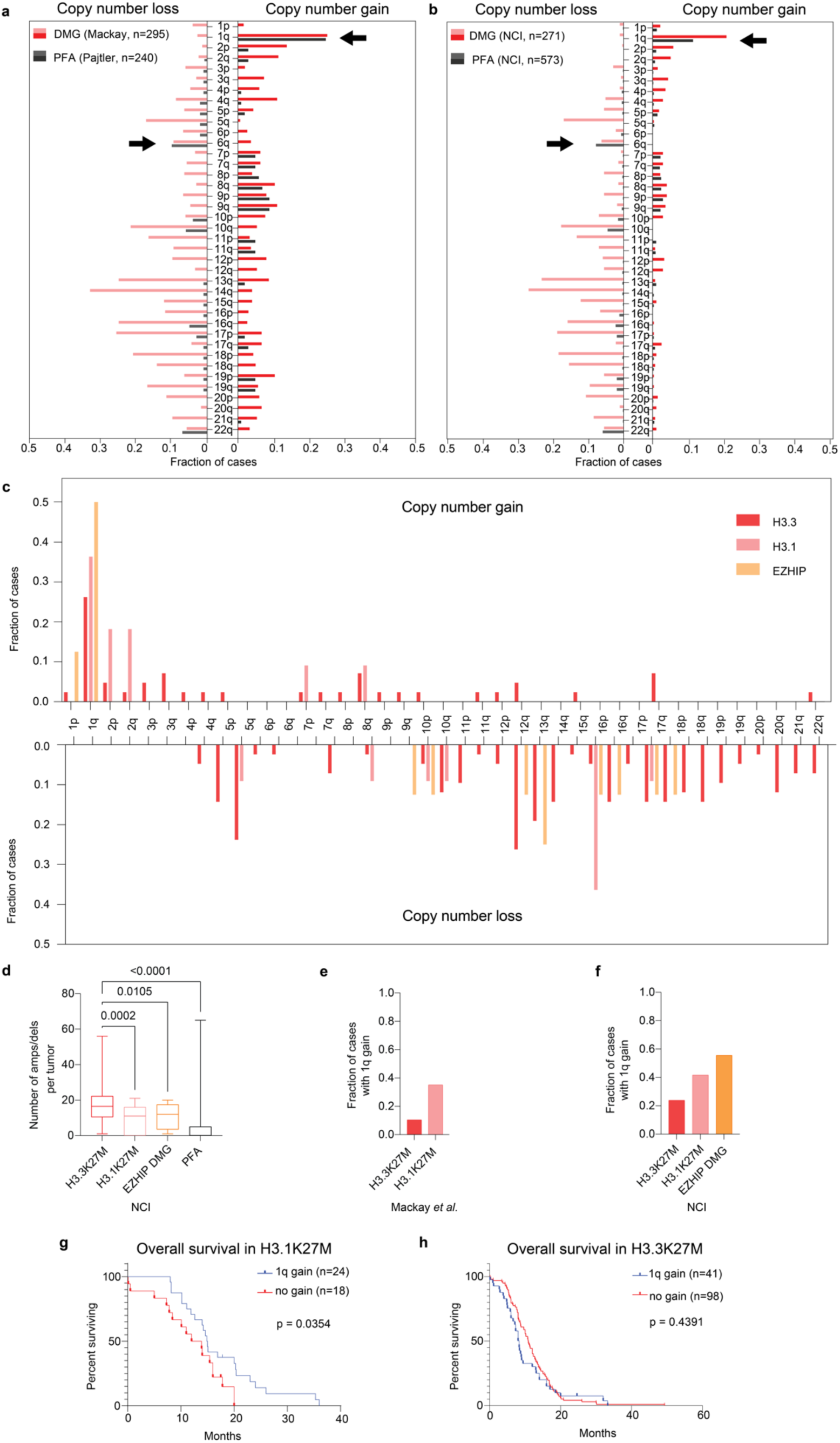
H3.1K27M and EZHIP-DMGs share key copy-number alterations with PFAs. **a** Copy number profiles (copy number loss left panel, copy number gain right panel) of H3K27M DMGs (Mackay *et al.*, 2017, n=295) [44] and PFAs (Pajtler *et al.*, 2015, n=240) [58] from published DNA methylation cohorts. Arrows indicate common shared recurrent alterations (more than 5% of tumors in all cohorts in **a** and **b** with known survival associations in PFAs. **b** Copy number profiles (copy number loss left panel, copy number gain right panel) of H3K27M DMGs (n=271) and PFAs (n=573) from the National Cancer Institute (NCI). Arm-level copy number gain/loss calls were made using GISTIC score cutoffs of +/-0.3. X-axis = chromosomes and Y-axis = fraction of cases. **c** Frequency of chromosome arm-level copy number alterations (copy number gain top panel, copy number loss bottom panel) for H3K27-altered DMGs from **b** segregated by molecular alterations - H3.3 mutant (n=42), H3.1 mutant (n=11) and H3-wildtype (H3- WT), EZHIP-expressing (n=8). X-axis = fraction of cases and Y-axis = chromosomes. **d** Quantification of the frequency of amplifications (amps) and deletions (dels, Y-axis) in DMGs segregated by molecular alterations (X-axis) and in PFAs from the NCI cohort. Data analyzed by one-way ANOVA with 95% confidence intervals. H3.3 mutant (n=48), H3.1 mutant (n=12), and EZHIP (n=8). **e** Frequency of 1q gain (Y-axis) for subtypes of H3K27M DMG and PFAs from published datasets depicted in **a**. H3.3 mutant (n=245), H3.1 mutant (n=49), and PFA (n=240). DMG sample pHGG_META_0223 was excluded from as it is H3.2-mutant. **f** Frequency of 1q gain (Y-axis) for subtypes of H3K27-altered DMG and PFA ependymoma from the NCI cohort depicted in 1b. H3.3 mutant (n=42), H3.1 mutant (n=11), and EZHIP (n=8). **g** Overall survival (months) analysis of 1q gain in H3.1K27M DMGs with (n=24) or without (n=18) 1q gain. **h** Overall survival (months) analysis of 1q gain in H3.3K27M DMGs with (n=41) or without (n=98) 1q gain. Data in 1g-h analyzed using Kaplan-Meier with Log-Rank test with 95% confidence intervals.

Gain of 1q has previously been associated with worse overall survival outcome in PFAs in multiple studies [24, 48, 57, 66, 72]. We noted that 1q gain was also the most common recurrent arm-length gain in H3K27M DMGs; however, there was no significant difference in overall survival in H3K27M DMGs with or without 1q gain from the published cohorts (**Supplementary Fig. 1a**). Previous work had demonstrated differences in the copy-number landscapes based on the type of H3 mutation with H3.3- versus (vs.) H3.1- mutant tumors exhibiting increased genomic instability [44]. We examined copy number trends with respect to H3.3 vs. H3.1 status in the NCI cohort and evaluated how H3 wildtype (H3-WT) EZHIP-expressing DMGs compared to each histone subtype. We observed that H3.1-mutant and EZHIP-expressing DMGs had fewer arm-length copy-number events (**Fig. 1c**) and focal amplifications and deletions (**Fig. 1d**) compared to H3.3 mutant DMGs. Gain of 1q was observed at a higher frequency in H3.1 vs. H3.3K27M DMGs in both the Mackay *et al*. (H3.1=35.1% vs. H3.3= 10.5%; **Fig. 1e**) and the NCI cohort (H3.1=41.7% vs. H3.3= 23.9%; **Fig. 1f**). Notably, half of EZHIP expressing DMGs also exhibited 1q gain (**Fig. 1f**). Unexpectedly, 1q gain was associated with better overall survival in H3.1K27M tumors (**Fig. 1g**), but not in H3.3K27M-mutant tumors (**Fig. 1h**).

In addition to recurrent 1q gain, the other alterations observed in at least 5% of each cohort were losses of 6q, 10q, and 22q (**Fig. 1a-b**). Loss of 6q has recently been associated with a subset of PFAs with poor survival rates [5]. Assessment of the 6q loss in the DMG cohort demonstrated a trend toward worse survival in 6q loss tumors (**Supplementary Fig. 1b**). In H3K27M DMGs, there was no major difference in 6q status when comparing H3.1 and H3.3-mutant cohorts (**Supplementary Fig. 1c**). Given the genomic instability of DMGs in general, we asked if 6q alterations were associated with high arm-level alterations. In every cohort, tumors with 6q loss were associated with higher levels of arm-level losses across the genome. In contrast, the frequency of arm gains in these tumors was not significantly different from non-6q loss tumors in each cohort (**Supplementary Fig. 1d**).

### *ACVR1* expression is prognostic in PFAs and is associated with transcriptomic signatures observed in *ACVR1*-mutant DMGs

Due to our observed differences between H3.1 and H3.3 K27M DMGs and 1q status, we extended our comparative analysis to include other genomic alterations that segregate with mutant histone subtypes. Activating mutations in activin A receptor type 1 (*ACVR1*) occur more frequently in H3.1 vs. H3.3 DMGs [9, 13, 21, 44]. Concordant with our findings of 1q gain enrichment in H3.1 tumors, we observed increased frequency of *ACVR1* mutations in 1q gain tumors vs. non-1q gain tumors (**Fig. 2a**).

**Fig. 2:**
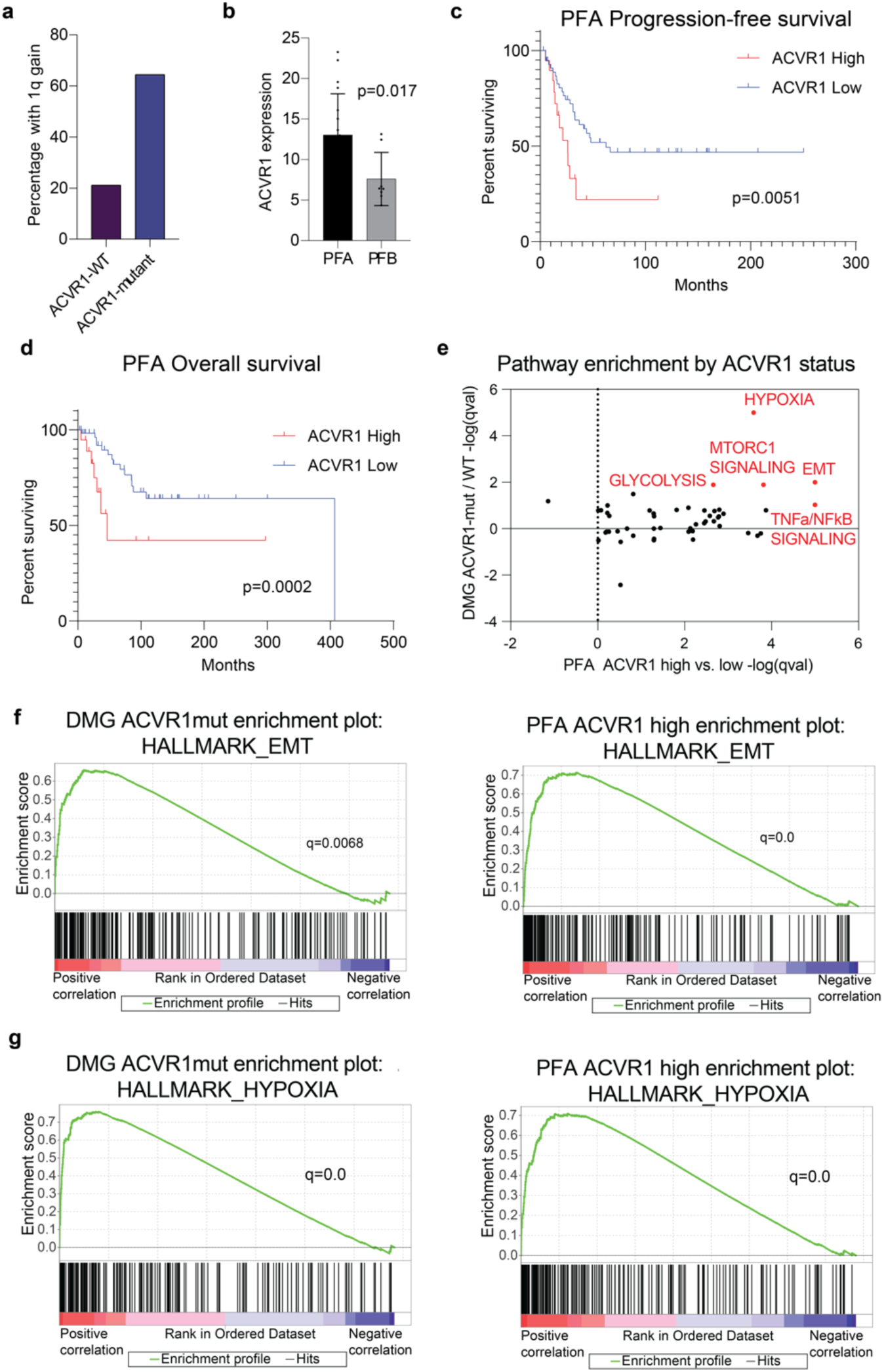
*ACVR1* expression is prognostic in PFAs and is associated with transcriptomic signatures observed in *ACVR1*-mutant DMGs. **a** Frequency of 1q gain (Y-axis) in *ACVR1*-mutant (n=31) vs. WT (n=137) H3K27M DMGs from Mackay *et al.* [44]. **b** *ACVR1* RNA expression levels (Y-axis) in PFAs (n=19) compared to PFBs (n=8) from Bayliss *et al*. 2016 [6]. Data analyzed by two-sided, unpaired, two-tailed *t* test. **c** Progression-free survival (months) analysis comparing PFAs with *ACVR1* high (the top quartile, n=19) vs. low (lower three quartiles, n=57) expression. **d** Overall survival (months) analysis comparing PFAs with *ACVR1* high (the top quartile, n=19) vs. low (lower three quartiles, n=57) expression. Data in **2c-d** analyzed using Kaplan-Meier, Log-Rank test with 95% confidence intervals. **e** Comparison of enriched pathways from differentially expressed genes in ACVR1 mutant vs. wildtype H3K27M DMGs (from 2a, Y-axis) and ACVR1 high vs. low PFAs (from 2d, X-axis). **f** Gene set enrichment analyses of the Hallmark epithelial-to-mesenchymal transition (EMT) pathway for genes ranked by differential expression in *ACVR1*-mutant DMGs or in high *ACVR1* expressing PFAs. **g** Gene set enrichment analyses of the Hallmark hypoxia pathway for genes ranked by differential expression in *ACVR1*-mutant DMGs or in high *ACVR1* expressing PFAs.

Recent work has identified rare mutations of *ACVR1* in PFA tumors [63]. These findings prompted us to investigate *ACVR1* in PF ependymomas. First, we observed increased expression of *ACVR1* in PFAs compared to group B posterior fossa ependymomas (PFBs), a subtype of PF ependymomas that lacks EZHIP expression [57] (**Fig. 2b**). We found tumors with the highest levels of *ACVR1* expression in the Pajtler *et al*. cohort to be associated with worse progression-free and overall survival outcomes (**Fig. 2c-d**). We next investigated whether ACVR1-mutant H3K27M DMGs and *ACVR1*-high PFAs shared transcriptomic signatures. We found that both tumors demonstrated enrichment of many ACVR1-mutant [31] and H3.1K27M-enriched [12] pathways including hypoxia/HIF1-α transcription factor network and epithelial-to-mesenchymal transition, as well as glycolysis, TNFa/ NFkB signaling, and mTORC1 signaling (**Fig. 2e-g**).

### H3K27me3 commonly enriched genes in PFAs and H3K27M DMGs exhibit heterogeneity corresponding to tumor anatomic location

We next focused our efforts on defining common epigenetic signatures in H3K27M DMGs and PFAs. Despite global reduction of H3K27me3, multiple studies have demonstrated that both tumors retain H3K27me3 at specific genomic sites [14, 27, 34, 51]. We performed a systematic comparison of the genes that retain H3K27me3 in H3K27M DMGs and PFAs to better understand this shared chromatin landscape.

H3K27me3 chromatin-immunoprecipitation with high throughput sequencing (ChIP-seq) profiles generated by our group [6] and others [8, 27, 43] was analyzed. We examined H3K27me3 enrichment in nine PFA ependymomas and six DMGs to define genes with commonly methylated promoters and gene bodies between the two tumors. We identified 551 genes with H3K27me3 retained in both tumor sets (**Fig. 3a**). Because H3K27me3 is a repressive mark, the extent of repression was examined by analyzing standard deviation values of gene expression for each of these 551 genes within both H3K27M DMGs and PFAs. Analysis of bulk expression data from both tumor cohorts identified high variance in expression levels of several H3K27me3-associated genes in both PFAs and H3K27M DMGs with greatest variability in homeobox genes including *HOXA2*, *HOXA4,* and *LHX2*. (**Fig. 3b**). Examination of genomic H3K27me3 at *HOX* gene clusters in PFAs, H3K27M DMGs and patient-derived H3K27M-DMG cell lines demonstrated distinct variations in enrichment (**Fig. 3c**). In both tumor sets, some samples showed H3K27me3 enrichment along the entire length of the *HOXA* and *HOXD* clusters, but others demonstrated variability in H3K27me3 at the *HOXA1* through *HOXA7* loci and the proximal HOXD locus (**Fig. 3c**).

**Fig. 3:**
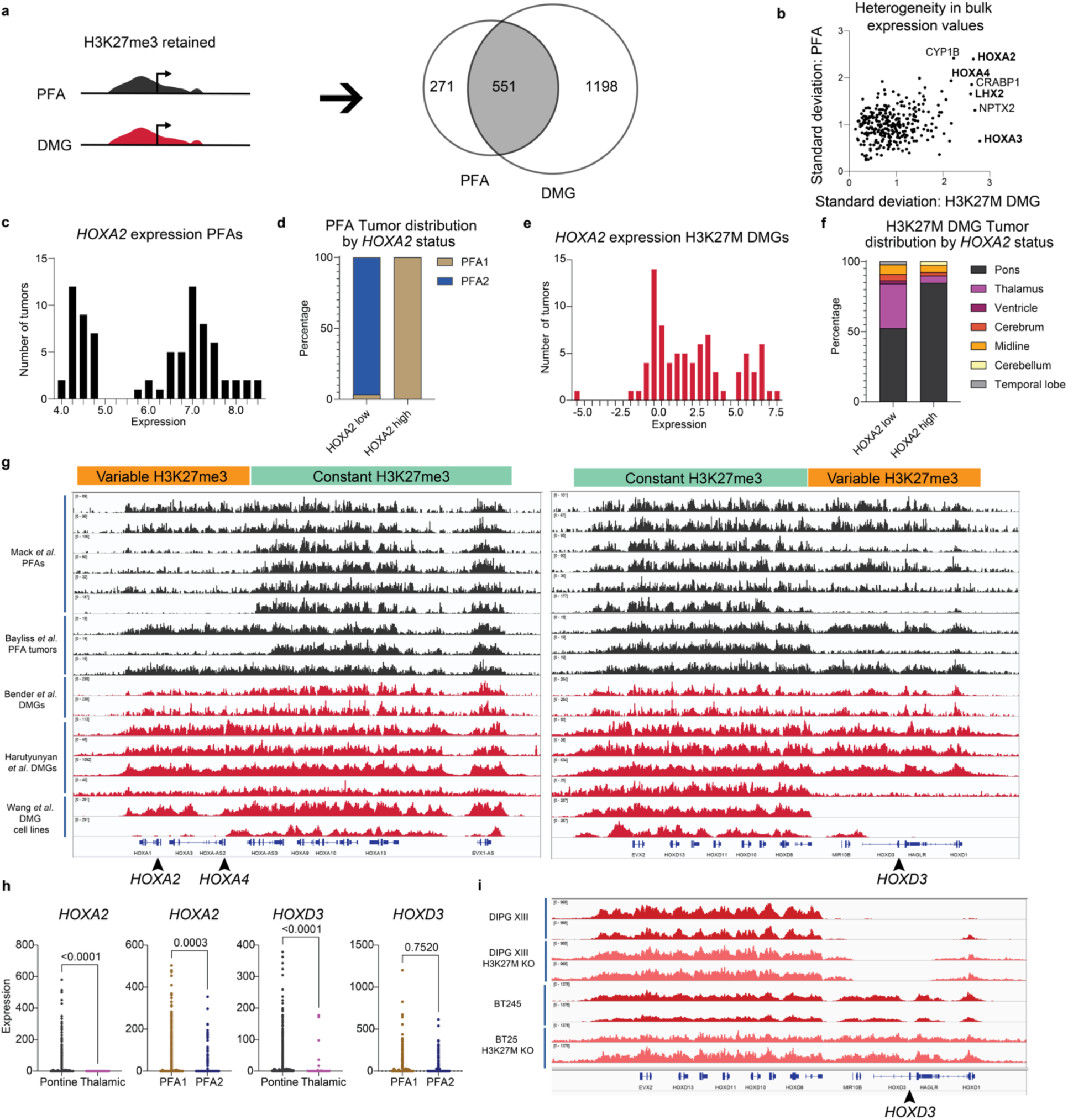
H3K27me3 commonly enriched genes in PFAs and H3K27M DMGs exhibit heterogeneity corresponding to tumor anatomic location. **a** Workflow of H3K27me3 ChIP-seq analysis. Genes with promoter (2kb upstream) or gene-body H3K27me3 peaks in more than two-thirds of PFAs (≥7/9 samples) and H3K27M DMGs (≥5/6) were identified as H3K27me3-retaining genes in each tumor type. The resulting gene sets were then compared to identify shared and tumor-specific H3K27me3-retaining genes. **b** Heterogeneity in expression of H3K27me3-retaining genes was identified by plotting the standard deviation in expression of each gene identified from 3a in H3K27M DMGs (X-axis) and PFAs (Y-axis). Genes, including *HOXD3*, for which expression data was not available in both bulk expression data sets were excluded from plotting. **c** H3K27me3 ChIP-seq tracks for PFAs (black, n=9, from Mack *et al*. [43], and Bayliss *et al*. [6]) and H3K27M DMGs (red, middle, n=6, from Bender *et al*. [8] and Harutyunyan *et al*. [27]) tumors and patient-derived H3K27M cell lines (red, bottom, n=2 from Wang *et al*. [74]) at the HOXA and HOXD clusters. Positions of proximal *HOX* genes associated with variation in expression (*HOXA2, HOXA4, HOXD3*) are indicated with arrowheads. Regions identified with differential patterns in H3K27me3 enrichment per tumor are labeled as variable (orange) and constant (green). Anatomic site for DMGs and PFA1/2 status were not available for tumor or cell-line samples. **d** Histogram of *HOXA2* expression values per tumor (X-axis = binned expression scores, Y-axis = number of tumors) in PFAs. **e** Comparison of *HOXA2* high (expression < 5 units) vs. low (expression > 5.5 units) groups based on the local minimum between modes in DNA-methylation defined, anatomically distinct PFA1 (n=49) and PFA2 (n=29) subtypes of PFAs. **f** Histogram of *HOXA2* expression values per tumor (X-axis = binned expression scores, Y-axis = number of tumors) in H3K27M DMGs. **g** Comparison of *HOXA2* high (expression < 4.5 units) vs. low (expression > 4.5 units) groups based on the local minimum between modes with anatomic location of H3K27M DMGs (pons=65; thalamus=26; midline not otherwise specified (NOS)=5; cerebellum=1, and other=2). **h** Expression patterns of *HOXA2* and *HOXD3* in single-cell RNA-sequencing (scRNA-seq) experiments (Filbin *et al.* and Gojo *et al*.) [20, 25]. Tumors were grouped by anatomic site (pons vs. thalamus for H3K27M DMG) or methylation-based subtype (PFA1 vs PFA2 for PFAs). **i** H3K27me3 ChIP-seq tracks at the HOXD locus from isogenic patient-derived H3K27M DIPG XIII and BT425 cell lines with or without H3K27M-knockdown.

*HOXA2*, *HOXA3,* and *HOXA4* have been identified as transcriptional markers that differentiate PFA1 and PFA2 [57]. Pajtler *et al*. proposed that the PFA1 tumors overexpress HOXA-family genes and hypothesized distinct anatomical origins PFA1 vs. PFA2 ependymomas [57]. Assessment of *HOXA2* bulk expression revealed a bimodal expression pattern in PFAs (**Fig. 3d**). As expected, segregation of *HOXA2* high vs. low PFAs occurred almost completely along with PFA1 vs. PFA2 status with majority of *HOXA2* low tumors corresponding to PFA2 subgroup (**Fig. 3e**). H3K27M DMGs similarly exhibited bimodal *HOXA2-*high vs. low distribution (**Fig. 3f**). Moreover, *HOXA2* expression levels revealed similar anatomic segregation with majority of thalamic tumors demonstrating low HOXA2 expression (**Fig. 3g**). *HOXA4* expression demonstrated a similar, though less striking pattern (**Supplementary Figs. 2a-d**). *HOXD3,* for which expression data in PFAs was unavailable, also followed a similar bimodal distribution (**Supplementary Fig. 2e**) and segregation with pontine tumors in *HOXD3*-high H3K27M DMGs (**Supplementary Fig. 2f**). Evidence of *HOX* family expression in single-cell data from DMGs and PFAs [20, 25] further supported this anatomic pattern in expression. Tumors derived from the pons and PFA1 showed higher *HOXA2* and *HOXA4* expression, while *HOXD3* expression was elevated in pontine vs. thalamic tumors but was not significantly different between PFA1 and PFA2 tumors (**Figs. 3h and Supplementary Fig. 2g**).

Differential H3K27me3 enrichment at these loci could be due to varying effects of PRC2 inhibition by the H3K27M mutation. To test this hypothesis, we examined published ChIP-seq tracks from DIPG-XIII and BT245 H3K27M patient-derived cell lines with or without H3K27M knockdown [27]. In both cell lines, H3K27M knockdown did not alter H3K27me3 genomic distribution at these genomic loci (**Fig. 3i**), suggesting that differential H3K27me3 at this site is not dependent on H3K27M. Overall, both H3K27M DMGs and PFA ependymomas showed differences in H3K27me3 genomic distribution at specific gene loci consistent with gene expression differences based on tumor anatomic location.

### Enhancer signatures in PFAs and H3K27M DMGs show high expression in astrocyte-like tumor cells

Both H3K27M mutations and EZHIP overexpression lead to a global increase in H3K27 acetylation (H3K27ac), a mark associated with active transcription [36, 53, 57, 59–61]. Moreover, H3K27M DMGs and PFAs exhibit unique H3K27ac marked enhancer and super enhancer profiles [37, 42, 52]. We combined information from gene lists generated by Mack *et al*. and Krug *et al*. and reanalyzed publicly available H3K27ac

ChIP-seq data from these studies to identify enhancers that are shared by, or unique to, each tumor (**Supplementary Fig. 3a**). H3K27ac-marked gene enhancers shared by both H3K27M DMGs and PFAs (n=533, **Supplementary Fig. 3**) were mainly associated with axonogenesis, axon guidance, and cell motility pathways, as well as differentiation of glial and oligodendrocyte lineages (**Fig. 4a**). DMG-specific enhancers mapped mainly to genes enriched for synaptic signaling (**Fig. 4b**), whereas PFA-specific enhancers were associated with genes enriched for extracellular organization and angiogenesis (**Fig. 4c**).

**Fig. 4.**
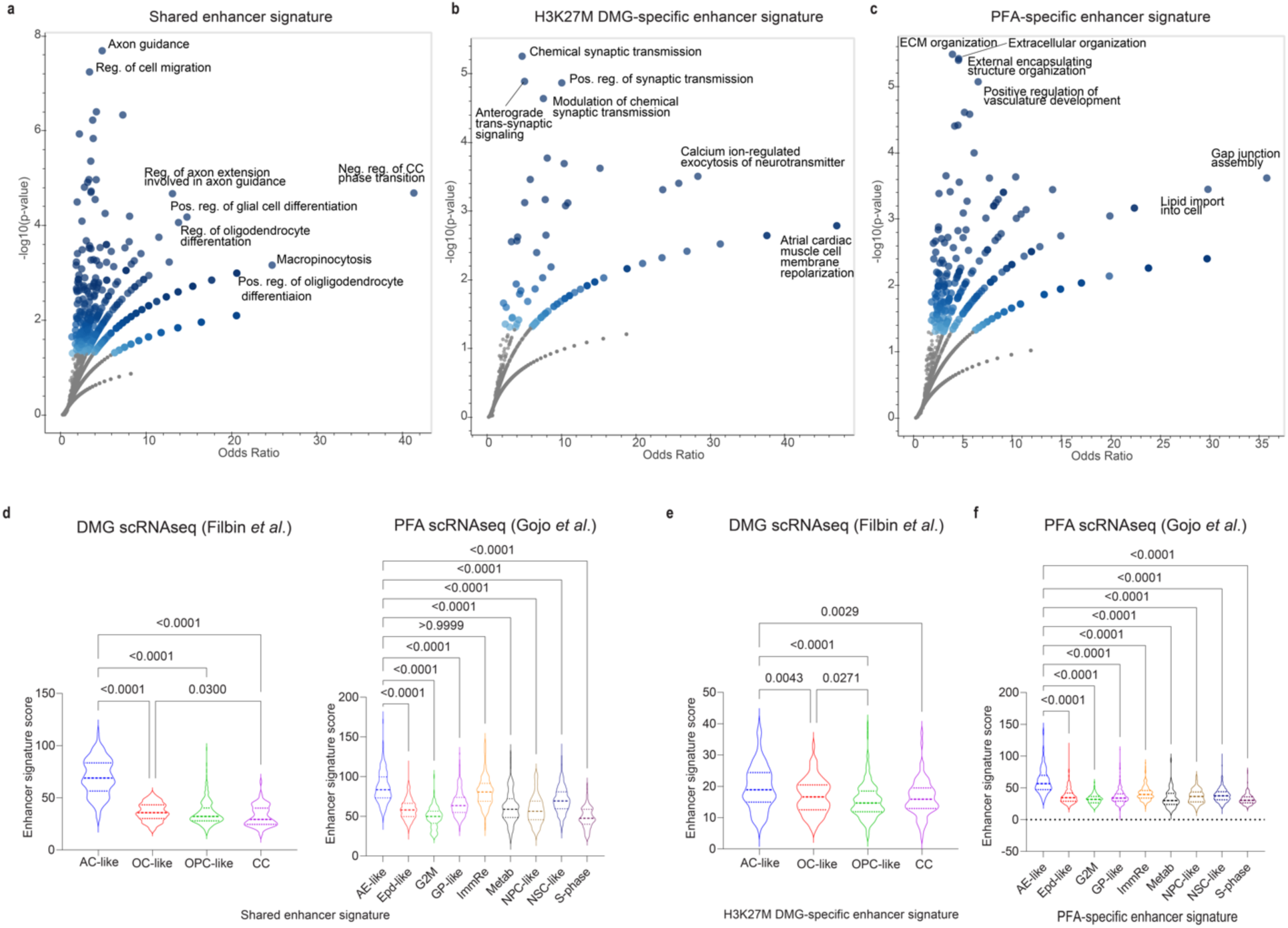
Enhancer signatures in PFAs and H3K27M DMGs show high expression in astrocyte-like tumor cells. **a** Enriched gene ontology biological process (GO BP) gene sets (Y-axis Negative Log10 p value, X-axis Odds ratio) for genes associated with enhancers shared by PFAs and H3K27M DMGs. Shading of points is influenced by p-value (darker shade with lower p-values) and number of overlapping points. **b** Enriched gene ontology biological process (GO BP) gene sets (Y-axis Negative Log10 p value, X-axis Odds ratio) for genes associated with enhancers unique to H3K27M DMGs. **c** Enriched gene ontology biological process (GO BP) gene sets for genes (Y-axis Negative Log10 p value, X-axis Odds ratio) associated with enhancers unique to PFA ependymomas. **d** Comparison of expression of the shared enhancer signature by cell type in H3K27M DMGs [20] and PFAs [25]. Signature scores per cell were calculated by taking the mean of expression values for the genes in the signature. **e** Expression of the H3K27M DMG-specific enhancer signature by DMG cell type. **f** Expression of the PFA-specific enhancer signature by DMG cell type. Data in 4d-f analyzed by non-parametric Kruskal-Wallis test and 95% confidence intervals. AC-like, Astrocyte-like; OC-like, Oligodendrocyte-like; OPC-like, Oligodendrocyte precursor-like; CC- cell cycle; AE-like, Astroependymal-like; Epd-like, Ependymal-like; G2M, G to M cell cycle related; GP-like, Glialprecursor-like; ImmRe, Immune related; Metab, Metabolic; NPC-like, Neuronal precursor-like; NSC-like, Neuronal stem cell-like; S-phase, S-phase cell cycle.

Due to the importance of enhancer activity in regulation of cell identity, we explored whether the enhancer signatures segregated with differentiation cell states within each tumor by assessing the single-cell RNA-seq datasets [20, 25]. Single-cell RNA- sequencing in H3K27M-DMGs and PFA ependymomas has demonstrated marked intratumoral heterogeneity [20, 25]. Both tumors contain malignant cells with varying expression of stem-cell-like, and more differentiated tumor cell signatures. H3K27M DMGs contain tumor cells bearing oligodendrocyte precursor (OPC)-like, oligodendrocyte (OC)-like and astrocyte (AC)-like cells, [20]. Similarly, PFAs contain less differentiated neuronal stem cell (NSC)-like, glial progenitor (GP)-like, and neuronal precursor cell (NPC)-like, as well as more differentiated ependymal (Epd)-like and astroependymal (AE)-like tumor cells [25]. Mapping the expression signatures onto tumor cell-types showed highest average expression of shared enhancer signature within astrocyte-like cells in H3K27M DMGs and astroependymal-like and immune- reactive PFA tumor cells (**Fig. 4d**). Evaluation of the tumor-specific enhancer signatures by single-cell type also showed increased expression in astrocytic-like lineages for both tumor types (**Fig. 4d-e**).

### H3K27me3-enriched *CRABP1* is highly expressed in progenitor populations of both tumors

In further interrogation of the single-cell datasets, we asked whether genes with retained H3K27me3 exhibited preferential expression in single-cell defined tumor cell populations. In both tumors, cells with progenitor-like states including OPC-like tumor cells in H3K27M-DMGs, and NPC-like tumor cells in PFA ependymomas, tended to have higher expression of shared genes (between H3K27M DMGs and PFAs) retained H3K27me3 signature (**Fig. 5a and b**).

**Fig. 5.**
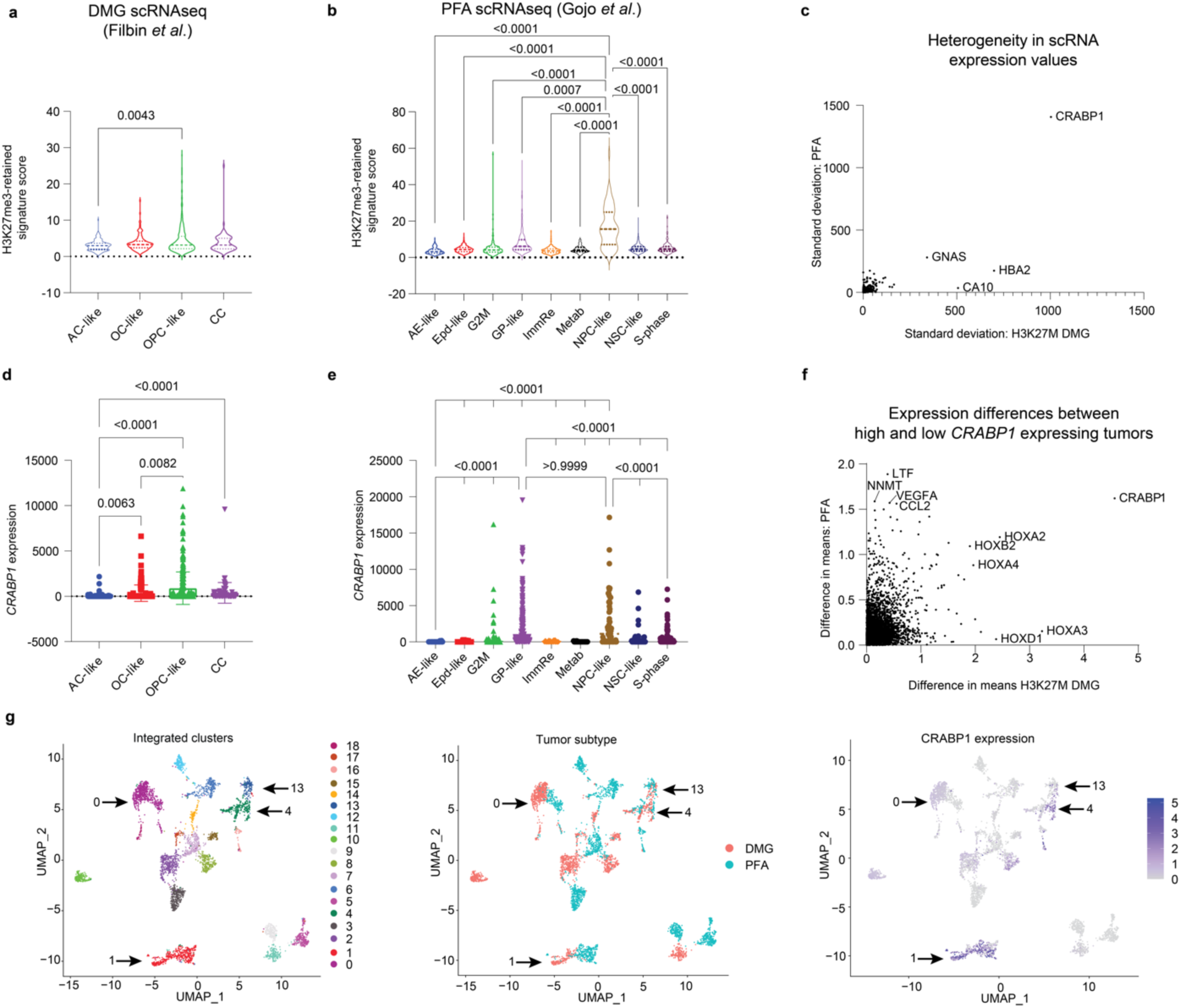
H3K27me3-enriched *CRABP1* is expressed highly in progenitor populations of both tumors. **a** Expression of the shared H3K27me3-retaining gene signature across cell-types in scRNA-seq data from H3K27M DMGs (Filbin *et al*.). Expression scores per cell were determined as the average expression of all genes in each cell. **b** Expression of the shared H3K27me3-retaining gene signature across cell-types in scRNA-seq data from PFA ependymomas (Gojo *et al*.). Only p-values for comparisons involving NPC-like cells are shown. **c** Comparison of the heterogeneity in expression of H3K27me3-retaining genes as measured by the overall standard deviation in expression of each gene from the H3K27me3 shared signature in H3K27M DMGs (X-axis) and PFA ependymomas (Y-axis). Genes for which expression data was not available in both bulk expression data sets were excluded from plotting. A total of 397 genes were plotted. **d** *CRABP1* expression values per cell from scRNA-seq data in H3K27M DMGs, grouped by cell type. **e** *CRABP1* expression values per cell from scRNA-seq data in PFAs, grouped by cell type. Only p-values for comparisons involving NPC-like and GP-like cells are shown. **f** Analysis of genes co-expressed with *CRABP1* in bulk RNA-seq datasets by comparison of difference in means for *CRABP1*-high and *CRABP1*-low tumors in H3K27M DMGs (X-axis) and PFA ependymomas (Y-axis). A split at the mean expression of *CRABP1* from the Mack *et* al. datset was used to delineate H3K27M DMG *CRABP1*-high (n=37) and *CRABP1*-low tumors (n=46). A split in the mean expression of the highest-level *CRABP1* probe from the Pajtler *et al*. dataset was used to delineate PFA *CRABP1*-high (n=22) and -low tumors (n=56). **g** (UMAP) embeddings of integrated scRNA-seq datasets of H3K27M DMG (Filbin) and PF ependymoma (Gojo). Left: identification of 20 distinct clusters. Middle: labeling by tumor of origin. Right: *CRABP1* expression across single cells with color scale minimum set to 0. Kruskal-Wallis test followed by multiple comparisons analysis were used to analyze data in Figs 5a-b and 5e-d.

To further assess heterogeneity, we analyzed genes with the greatest variability in expression (by comparing standard deviation values for each gene) across all tumor cell types in both tumors. Expression of cellular retinoic acid binding protein 1 (*CRABP1*) was a clear outlier for both tumor subtypes (**Fig. 5c**). Moreover, *CRABP1* was one of the H3K27me3 marked genes (**Supplementary Fig. 4a**) with the greatest variability in expression levels in the bulk datasets (**Fig. 3b**). *CRABP1* is frequently dysregulated in cancer [19, 40, 41, 71, 78]. *CRABP1* expression segregated strongly with OPC-like cells in DMGs (**Fig. 5d**) and both glial progenitor-like (GP-like) and neuronal precursor-like (NPC-like) in PFAs (**Fig. 5e**). We determined genes that correlate with *CRABP1* expression in bulk expression to find that the *HOX* genes were among the strongest correlates in both tumor datasets (**Fig. 5f**). However, *HOXA2* and *HOXD3* did not demonstrate any distinct patterns of expression by cell-type in DMGs, and the significantly different changes observed in PFAs were far less drastic (**Supplementary Figs. 4b-c**).

The scRNA-seq datasets were integrated to further explore similarities in expression patterns of H3K27M DMGs and PFAs. Integrated dimensional reduction analyses identified 20 clusters, four of which (clusters 0, 1, 4, and 13) included large populations of both PFA- and DMG-derived cells (Arrows, **Fig. 5g**). The top marker gene in the second-largest cluster’s (cluster 1) was *CRABP1* (**Fig. 5g, Supplementary Fig. 4d**). This cluster consisted primarily of progenitor and precursor cell states: OPC-like cells from the DMGs and GP-like cells from PFAs (**Supplementary Fig. 4e**). Overall, these data support a role for *CRABP1* in specific tumor cell-states in both tumors.

### Common H3K27me3 signatures mirror human hindbrain brain developmental patterns

H3K27M DMG and PFAs are proposed to arise from cells along developmental lineages in the hindbrain [20, 22, 25, 35, 73]. We first aimed to identify whether H3K27me3- associated genes exhibited specific patterns in normal human hindbrain development. We utilized published scRNA-seq dataset from fetal human cerebellum ranging from 9-21 weeks post-conception published by Aldinger *et al.* 2021 [1]. Overall, average expression of H3K27me3-retained genes increased in more differentiated populations of cells associated with potential cells-of-origin for DMGs and PFAs. Committed OPCs demonstrated increased signature expression compared to OPCs, and brainstem- choroid/ependymal cells had higher levels of expression than early brainstem cells (**Fig. 6a**). Cells of astrocytic lineage did not show a similar pattern (**Fig. 6a**). Because OPCs are the leading cell of origin for H3K27M DMGs [20, 52], we examined oligodendroglial lineages in more detail in a second single-cell dataset of human developing cerebellum (Sepp *et al.*, unpublished) [68]. H3K27me3 signatures were low in progenitor cells but increased along stages of OPC differentiation with highest expression in late-OPCs and then decreased in more differentiated committed OPCs and oligodendrocytes (**Fig. 6b**). Analysis of the astrocytic lineage cells did not yield a similar pattern (**Fig. 6c**).

**Fig. 6.**
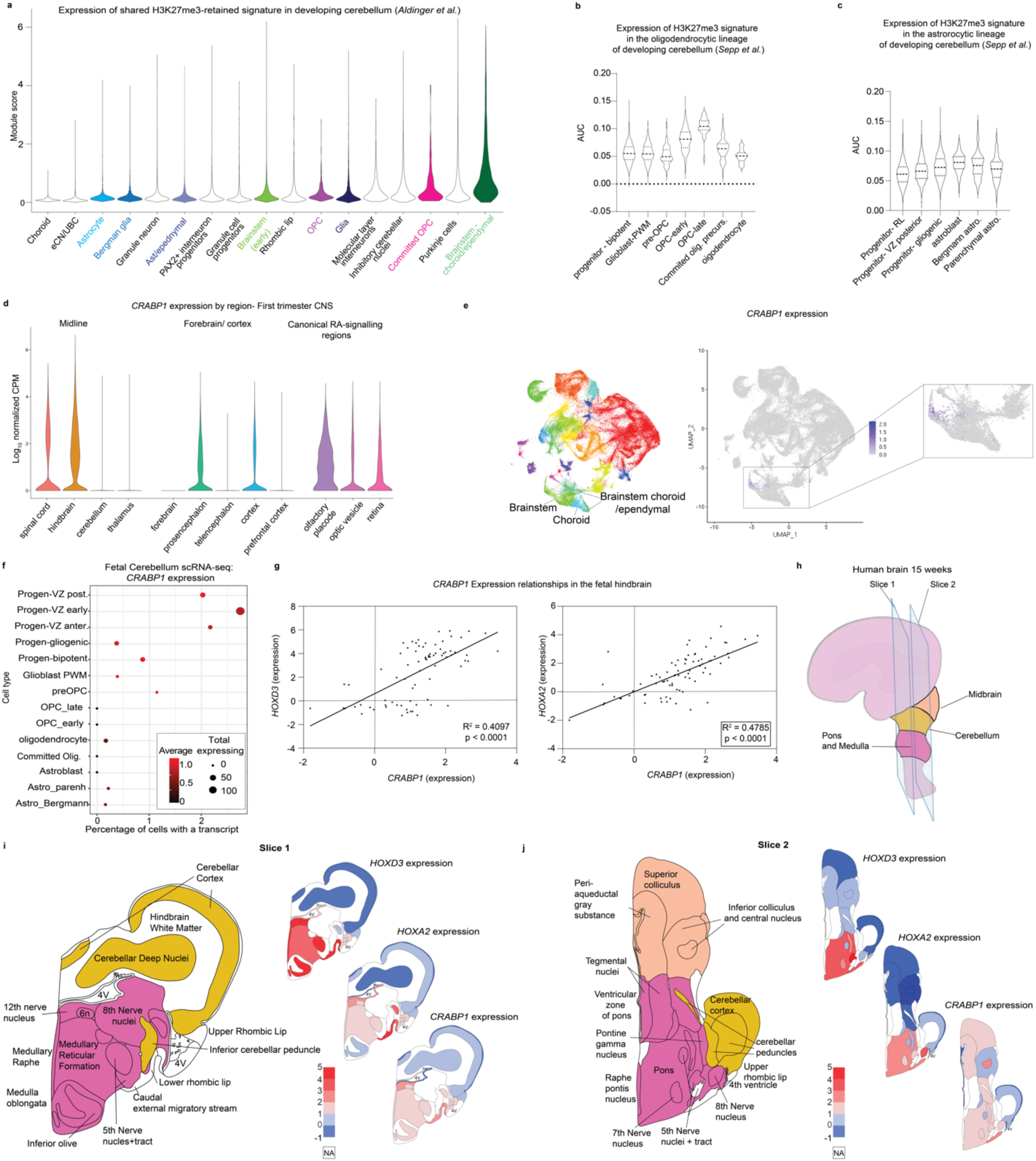
Common H3K27me3-retained signature mirror human hindbrain brain developmental patterns. **a** Expression of the shared DMG/PFA-derived H3K27me3 gene signature in scRNA-seq data of the developing human posterior fossa (Aldinger *et al*.) [1] grouped by cell-type. Highlighted groups indicate cell types pertinent to DMGs/PFAs. **b** H3K27me3 gene signature expression along the oligodendrocyte lineage in an independent scRNA-seq atlas of the developing posterior fossa (Sepp *et al*., unpublished) [68]. **c** H3K27me3 gene signature expression along the astrocytic lineage in the scRNA-seq atlas of the developing posterior fossa used in **b**. **d** First-trimester human brain scRNA-seq [18] expression patterns of *CRABP1* grouped by anatomic region. Left: midline structures, middle: forebrain and cortical structures, right: regions with canonical retinoic acid (RA) signaling. **e** *CRABP1* expression patterns in the developing cerebellum scRNA-seq atlas from **a**. Left: UMAP embedding colored by cluster, middle: *CRABP1* expression across all cells in UMAP embedding, right (inset): magnified depiction of *CRABP1* expression patterns in a subset of cells including brainstem and brainstem-derived choroidal/ependymal cells. **f** *CRABP1* expression patterns in astrocytic and oligodendrocyte lineages of the developing cerebellum scRNA-seq atlas from **b** and **c**. Early progenitors are grouped along the top of the Y-axis, then the more differentiated oligodendrocyte and the astrocytic lineages are each grouped below along the Y-axis. The percentage of cells expressing at least one *CRABP1* transcript (X-axis), the total number of cells expressing at least one *CRABP1* transcript (point size) and the average expression value (color) are depicted. **g** Microarray-based log2 expression levels from micro-dissected fetal hindbrain tissue at 15 and 16 weeks post-conception from BrainSpan’s Prenatal lateral microdissection (LMD) Microarray comparing *CRABP1* expression patterns to those of *HOXA2* and *HOXD3*. Data were analyzed with a simple linear regression and R-squared calculated with a goodness of fit test with 95% confidence intervals. **h** Schematic depicting sectioning of the 15-16 weeks post conception fetal brain utilized for depictions in **g**, **I**, and **j**. Early anatomic structures of the infratentorial brain are colored in peach (midbrain), yellow (cerebellum), and deep pink (pons and medulla). **i-j** Expression of *HOXD3*, *HOXA2*, and *CRABP1* expression heatmaps at 15 and 16 weeks post conception infratentorial brain at slices 1 (**i**) and slices 2 (**j**) from **h**. Left: schematic of brain sub-structures colored by overarching structure corresponding to key in **h**. Right, from top to bottom heatmaps of *HOXD3*, *HOXA2*, and *CRABP1* expression in corresponding areas. Illustration adapted from reference figure from the Allen Brain Atlas.

Given its notable expression pattern in the single-cell tumor datasets (**Fig. 5**), we examined *CRABP1* expression in the developing hindbrain utilizing a scRNA-seq dataset generated from human brain samples from the first trimester [18]. Carnegie stages are used to define period of human brain development [56]. In the hindbrain, we noted a peak of *CRABP1* expression in stage 19, which corresponds to the peak in expression of *H3-3A*, the histone variant most commonly mutated in DMGs (**Supplementary Fig. 5a**). With respect to anatomic location, *CRABP1* expression was higher in the spinal cord and hindbrain vs. thalamus and cerebellum (**Fig. 6d**). Known areas of high retinoic acid (RA)-signaling in development, including the olfactory placode and optic pathways demonstrated enrichment of *CRABP1* expression, whereas expression in forebrain/cortical-related structures was variable (**Fig. 6d**). Similarly, *CRABP1* expression was restricted to a subset of brainstem cells in the Aldinger *et al*. developing cerebellum atlas (**Fig. 6e**). At a cellular level, *CRABP1* expression was highest in the progenitor cell types compared to their more differentiated oligodendrocytic and astrocytic cells (**Fig. 6f**).

Data from mouse development have previously identified *CRABP1* expression as specific to rhombomeres 4-6 (r4-6) [45]. The r4-6 segments give rise to the caudal portion of the basilar pons (r4) and retropontine structures (r5-6) [75, 76]. Given this spatial patterning, the Allen Brain Atlas’ Developing Human Transcriptome was examined to determine patterns of *CRABP1* expression [2, 50]. Within hindbrain structures CRABP1 expression strongly correlated with *HOX* genes including *HOXA2* and *HOXD3* (**Fig. 6g**). Moreover, HOXD3, HOXA2 and CRABP1 expression in the developing human hindbrain at 15-weeks gestation was mainly restricted to developing human pontine and medullary structures (**Figs. 6h-j**).

## Discussion

While EZHIP-expressing PFAs and H3K27-altered DMGs (including H3.3 and H3.1K27M gliomas and EZHIP positive DMGs) have distinct natural histories and histologic features, evidence of their shared molecular features continues to mount as we profile broader samples of these tumor types. Furthermore, both tumors most commonly arise in neighboring hindbrain/posterior fossa-derived structures. This pattern suggests that developmental lineages in this region possess susceptibility to alterations that impede canonical PRC2 activity and therefore regulation of H3K27me3 state. Leveraging the existing molecular knowledge about each of these tumor types, we exploited genomic, bulk transcriptomic, scRNA-seq, and epigenomic profiles of both cancers to elucidate shared features that may improve our understanding of tumor biology.

Assessment of the genomic landscape of these tumors demonstrated that H3.1K27M DMGs showed greater copy number similarity with EZHIP-expressing DMGs than did H3.3K27M tumors, with more quiescent landscapes and higher frequencies of 1q gain. These profiles of relatively lower genomic instability may point toward greater commonalities between PFAs and H3.1K27M vs. H3.3K27M DMGs. Consistent with this pattern, we identified high *ACVR1*—a gene frequently altered in H3.1-mutant DMGs— expression to be associated with: (a) worse survival in PFAs, and (b) upregulation of pathways that were also enriched in ACVR1-mutated gliomas.

We additionally characterized common and unique features of H3K27M DMG and PFA chromatin landscapes based on patterns of H3K27me3/ac genomic deposition. The two tumors exhibited a considerable overlap in both H3K27me3-marked and H3K27ac- enhancer-associated genes. Exploration of the transcriptional states associated with these gene sets demonstrated inter- and intra-tumoral heterogeneity of expression patterns. Common enhancer-associated genes enriched for cell motility, and oligodendrocyte and glial differentiation pathways. Moreover, scRNA-seq demonstrated the highest levels of expression of these genes in astrocytic lineage cell types, especially in comparison to progenitor cell types. These patterns highlight the unique insights single-cell resolution transcriptomics can provide and illustrate how tumor-specific enhancers generated from bulk-sequencing methods may not fully reflect the regulatory landscapes of the tumors.

Interrogation of genes associated with residual H3K27me3 also yielded important insights. The sensitivity of H3K27M DMGs and PFAs to EZH2 inhibition [43, 49, 51] points to a potential role for H3K27me3 in maintaining tumor cell proliferative capacity. Data from developing cerebellar atlases provide some support for this model in that upregulation of genes within these signatures is seen along oligodendrocyte lineages and in brainstem choroid/ependymal cells. Furthermore, we noted distinct expression profiles of proximal vs. distal *HOX* genes, which correlated with inferred anatomic origins of tumor, and these genes demonstrate distinct anterior-posterior boundaries of expression in the developing human hindbrain. These data suggest that developmental epigenetic profiles and origins of PFAs and H3K27M DMGs may influence the H3K27me3 landscape and gene expression profile in these tumors.

*CRABP1,* like the *HOX* genes, has been shown to have highly specific expression patterns in mouse nervous system development [45]. In both PFAs and DMGs, its expression correlated with specific tumor cell types, with an enrichment among progenitor-like cells. While its precise role remains unclear, *CRABP1* has been proposed to be a critical regulator of retinoic acid signaling, preventing canonical RA-induced transcriptional programs [54]. This role is of particular interest due to retinoic acid’s function in hindbrain patterning [7, 23, 30, 47, 55] and the previous identification of the retinoic acid binding protein RXRA as the top ranked super-enhancer associated gene in H3K27M DMGs [52]. Together these data suggest that *CRABP1* may be associated with progenitor cells in early stages of the developing hindbrain that may be deregulated in both PFAs and H3K27M DMGs. Further investigation is needed to elucidate whether *CRABP1* contributes to maintenance of progenitor-type states in the presence of retinoids during development.

Overall, our studies further define molecular commonalities in the biology between DMGs and PFAs that express H3K27M and EZHIP and highlight similar patterns in the heterogeneity among tumors of each class. Moreover, some of these common epigenetic signatures showed specific patterns in human hindbrain development supporting the overall hypothesis that hindbrain developmentally and epigenetic deregulated molecular mechanisms may drive the biology of both tumors.

## Declarations

### Ethics approval and consent to participate

No consent to participate was required for the present study.

### Consent for publication

All authors consented to publication.

### Availability of data and material

Data were obtained as described in the Methods section.

### Funding and Acknowledgements

The work was supported in part by grants from National Institute of Neurological Disorders and Stroke (NINDS) R01NS110572 (SV), National Cancer Institute (NCI) R01ACA261926 (SV), NCI R37CA214955-01A1 (AR), NCI P30CA046592 (AR), Chad Tough Defeat DIPG foundation (SV), Alex Lemonade Stand Foundation (SV), the Hyundai Hope on Wheel Foundation (SV), and the Rogel Cancer Center PIBS Graduate Student Scholarship (MP).

### Authors’ contributions

MP and SV conceptualized the project and wrote the manuscript. MP, DP, PRN, PKJ, LJ, BE, MC, and SV performed formal analyses. MP, PRN, PKJ, and SV created visualizations of the data. AR, KA, SP, MF, AMC, AB, and SV supervised the analyses and provided valuable feedback. All authors reviewed, edited, and approved the manuscript.

---

**Supplementary Fig. 1:**
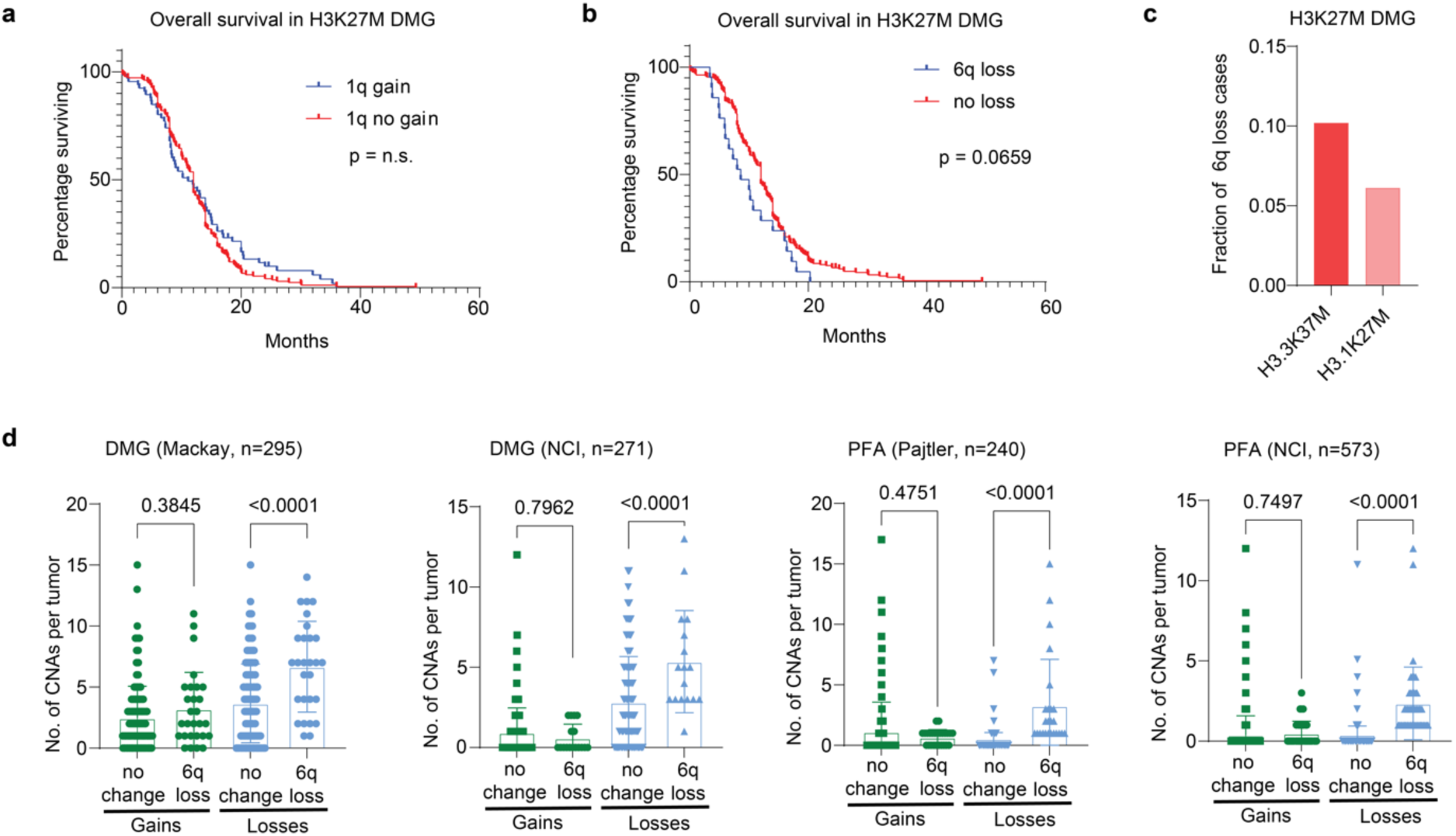
H3.1K27M and EZHIP-DMGs share key copy-number alterations with PFAs. **a** Overall survival (months) analysis of 1q gain in H3K27M DMGs with (n=66) or without (n=178) 1q gain. **b** Overall survival (months) analysis of 6q gain in HK27M DMGs with (n=21) or without (n=216) 6q loss. Data in S1a-b analyzed using Kaplan-Meier, Log-Rank test with 95% confidence intervals. **c** Frequency of 6q loss (Y-axis) for subtypes of H3K27M DMG from published datasets depicted in 1a. H3.3 mutant (n=245) and H3.1 mutant (n=49z). **d** Number of copy number alterations (CNAs, Y-axis) in DMGs and PFAs. All four cohorts were segregated by tumors overall chromosomal gains (green) with (n=28- Mackay, 17-NCI:DMG, 25-Pajtler, 45-NCI:PFA) or without 6q loss (n=267-Mackay, 254- NCI:DMG, 215-Pajtler, 528-NCI:PFA), or overall chromosomal losses (blue). Data analyzed by two-sided, unpaired, two-tailed *t* test.

**Supplementary Fig. 2:**
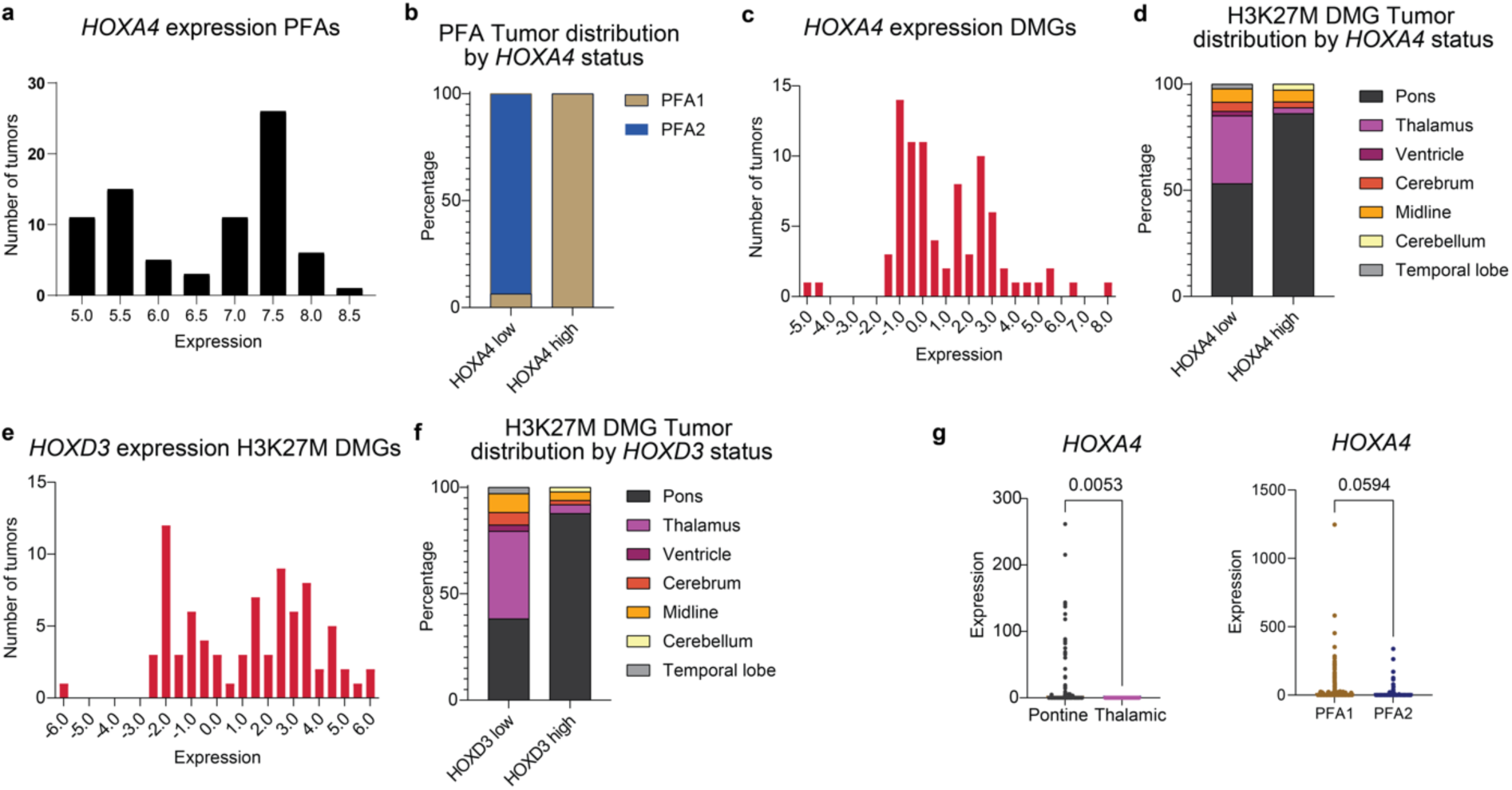
H3K27me3 commonly enriched genes in PFAs and H3K27M DMGs exhibit heterogeneity corresponding to tumor anatomic location. **a** Histogram of *HOXA4* expression values per tumor (X-axis = binned expression scores, Y-axis = number of tumors) in PFAs. **b** Comparison of *HOXA4* high (expression < 6 units) vs. low (expression > 6 units) groups based on the local minimum between modes in DNA-methylation defined, anatomically distinct PFA1 (n=49) and PFA2 (n=29) subtypes of PFAs. **c** Histogram of *HOXA4* expression values per tumor (X-axis = binned expression scores, Y-axis = number of tumors) in H3K27M DMGs. **d** Comparison of *HOXA4* high (expression < 1 unit) vs. low (expression > 1 unit) groups based on the local minimum between modes with anatomic location of H3K27M DMGs (pons=65; thalamus=26; midline not otherwise specified (NOS)=5; cerebellum=1, and other =2). **e** Histogram of *HOXD3* expression values per tumor (X-axis = number of values, Y-axis = bin center) in H3K27M DMGs. Note that *HOXD3* expression values were not available for PFAs. **f** Comparison of *HOXD3* high (expression > 0.5 units) vs. low (expression < 0.5 units) groups based on the local minimum between modes with anatomic location of H3K27M DMGs (pons=65; thalamus=26; midline not otherwise specified (NOS)=5; cerebellum=1, and other=2). Note that *HOXD3* expression values were not available for PFAs. **g** Expression patterns of *HOXA4* in single-cell RNA-sequencing (scRNA-seq) experiments (Filbin *et al.* and Gojo *et al*.) [20, 25]. Tumors were grouped by anatomic site (pons vs. thalamus for H3K27M DMG) or methylation-based subtype (PFA1 vs PFA2 for PFAs).

**Supplementary Fig. 3:**
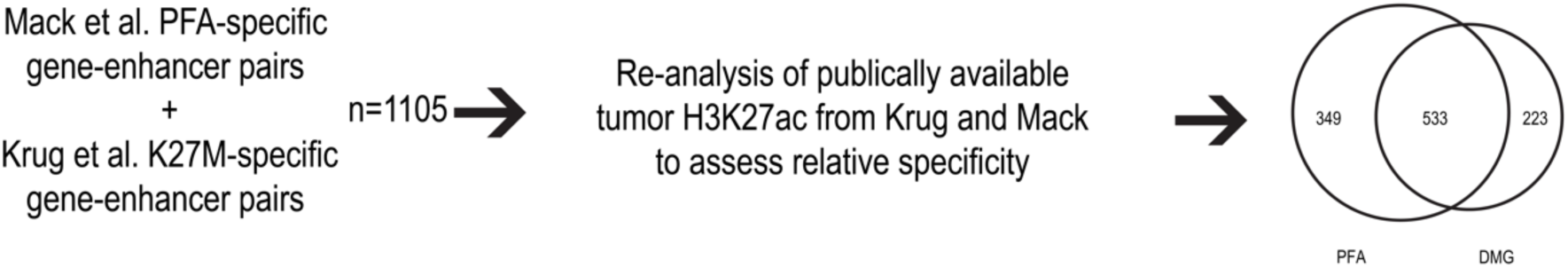
Enhancer signatures in PFAs and H3K27M DMGs show high expression in astrocyte-like tumor cells. **a** Venn diagram depicting the construction of the shared and tumor-specific enhancer signatures.

**Supplementary Fig. 4.**
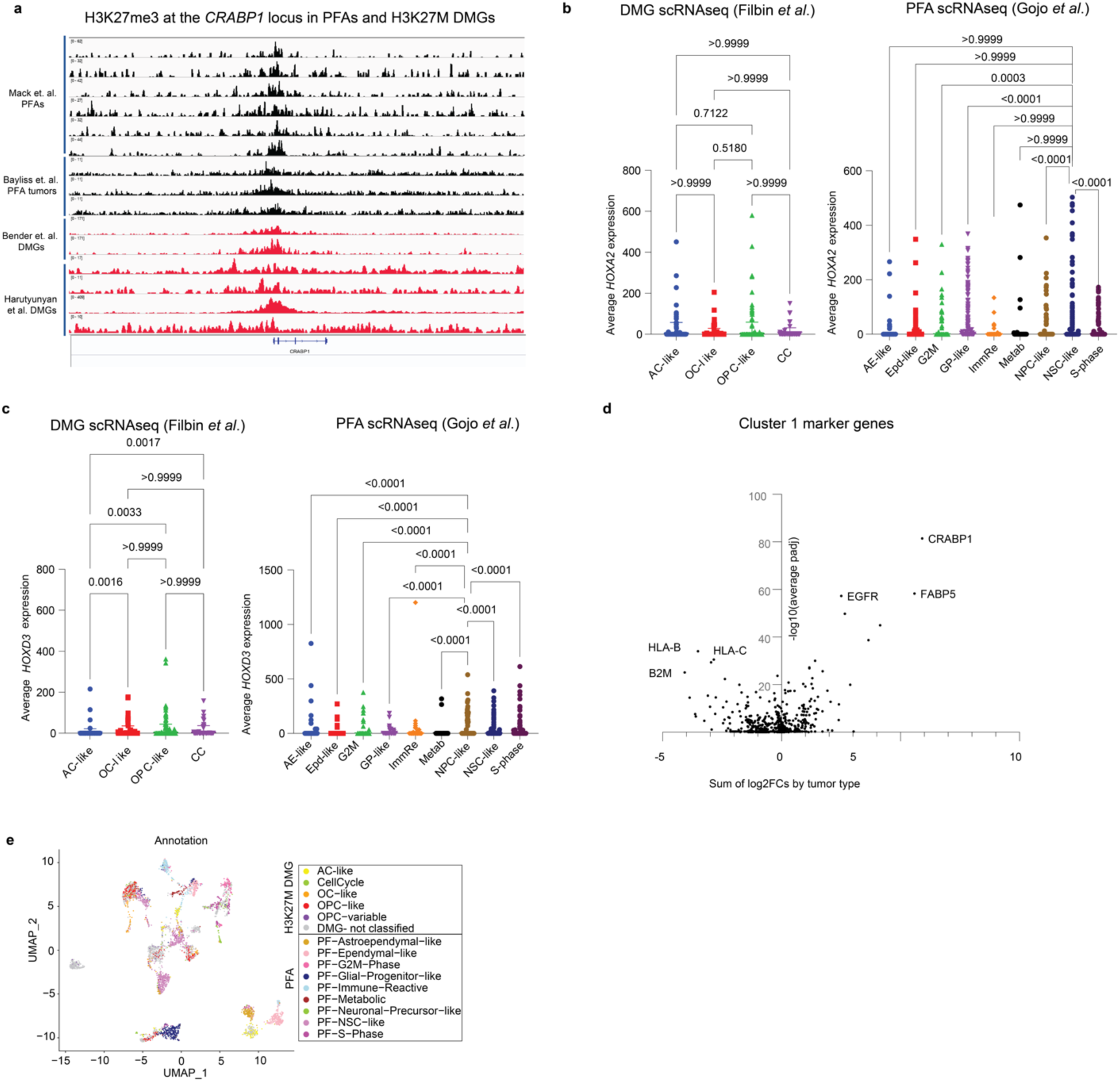
H3K27me3 enriched *CRABP1* is expressed highly in progenitor populations of both tumors. **a** H3K27me3 ChIP-seq tracks at the *CRABP1* locus for PFA (black, n=9, from Mack *et al*. [43], and Bayliss *et al*. [6]) and H3K27M DMG (red, n=6, from Bender *et al*. [8] and Harutyunyan *et al*. [27]) tumors. **b** scRNA-seq expression of *HOXA2* grouped by cell-type in PFA [25] and DMG [20] datasets. **c** scRNA-seq expression of *HOXD3* grouped by cell-type in PFA [25] and DMG [20] datasets. **d** Volcano plot of Seurat conserved markers of Cluster 1 in the integrated H3K27M DMG/PFA scRNA-seq analysis from Fig. 4g (Y-axis = -log10-normalized average adjusted p-value across the two tumor sets, X-axis = sum of log2-transformed fold changes between Cluster 1 and all other clusters for the two tumor sets. Kruskal-Wallis test followed by multiple comparisons analysis were used to analyze data in Figs S4b-c.

**Supplementary Fig. 5.**
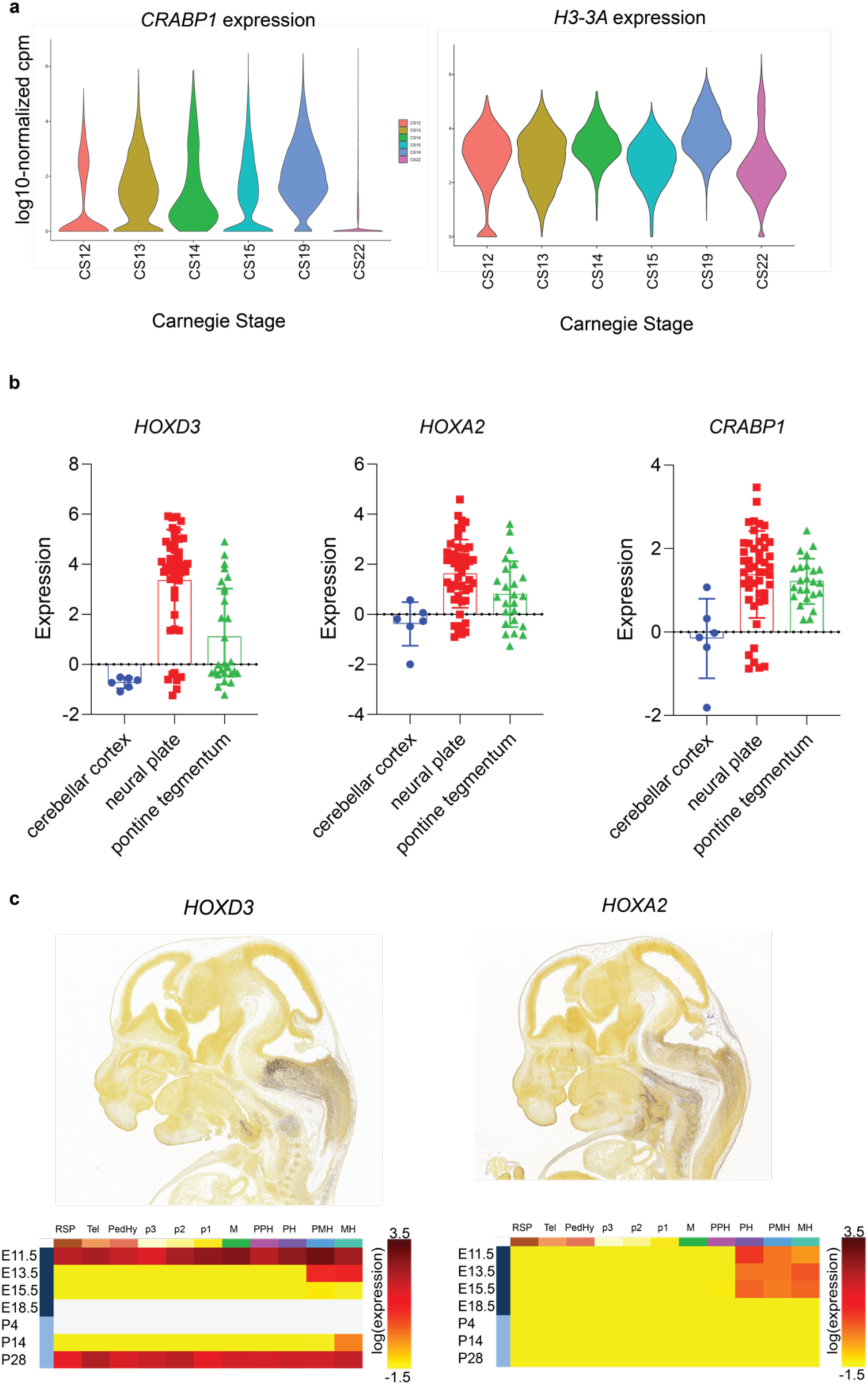
Common H3K27me3 signatures mirror human hindbrain brain developmental patterns. **a** First-trimester hindbrain *CRABP1* and *H3-3A* expression by Carnegie Stage (CS) of development [56] **b** Expression patterns of *CRABP1*, *HOXA2*, and *HOXD3* at 15 and 16 weeks post- conception in the BrainSpan prenatal LMD microarray dataset grouped by hindbrain substructures [2, 50]. **c** *HOXA2* and *HOXD3* expression patterns of the developing mouse from the Allen Brain Atlas. Top: *in situ* hybridization (ISH) staining for *HOXA2* and *HOXD3* in embryonic day 13.5 (E13.5). Bottom: Expression patterns of *HOXA2* and *HOXD3* across space and time in the developing mouse brain: RSP, rostral secondary prosencephalon; Tel, telencephalic vesicle; PedHy, peduncular hypothalamus; p3, prosomere 3; p2, prosomere 2; p1, prosomere 1; M, midbrain; PPH, prepontine hindbrain; PH, pontine hindbrain; PMH, pontomedullary hindbrain; MH, medullary hindbrain. Image credit: Allen Institute for Brain Science: https://developingmouse.brain-map.org/experiment/siv?id=100093342&imageId=101522680&initImage=ish, (*HOXD3*ISH) https://developingmouse.brain-map.org/gene/show/15209 (*HOXD3* heatmap), https://developingmouse.brain-map.org/experiment/siv?id=100030618&imageId=100662735&initImage=ish (*HOXA2*ISH) https://developingmouse.brain-map.org/gene/show/15174 (*HOXA2* heatmap).

